# NNAL, a major metabolite of tobacco-specific carcinogen NNK, promotes lung cancer progression through deactivating LKB1 in an isomer-dependent manner

**DOI:** 10.1101/2021.06.15.448557

**Authors:** Tengfei Bian, Yuzhi Wang, Jordy F. Botello, Qi Hu, Yunhan Jiang, Adriana Zingone, Pedro A. Corral, F. Zahra Aly, Yougen Wu, Bríd M. Ryan, Lingtao Jin, Chengguo Xing

**Affiliations:** Department of Medicinal Chemistry, University of Florida, 1345 Center Drive, Gainesville, FL 32610, USA; Department of Anatomy and Cell biology, University of Florida, Gainesville, FL 32610, USA; Laboratory of Human Carcinogenesis, Center for Cancer Research, National Cancer Institute, National Institutes of Health, Bethesda, MD 20892, USA; Department of Pathology, Immunology and Laboratory Medicine, University of Florida, 1345 Center Drive, Gainesville, FL 32610, USA; College of Tropical Agriculture and Forestry, Hainan University, 58 Renmin Avenue, Haikou 570228, China

**Author notes:** These two authors contributed equally to this paper (Tengfei Bian, Yuzhi Wang).

## Abstract

Smoking is associated with worse clinical outcomes for lung cancer patients. Cell-based studies suggest that NNK (a tobacco specific carcinogen) promotes lung cancer progression. Given its short half-life, the physiological relevance of these *in vitro* results remains elusive. NNAL, a major metabolite of NNK with a similar structure, a chiral center, and a longer half-life, has never been evaluated in cancer cells. In this study, we characterized the effect of NNAL and its enantiomers on cancer progression among a panel of NSCLC cell lines and explored the associated mechanisms. We found that (R)-NNAL promotes cell proliferation, enhances migration, and induces drug resistance while (S)-NNAL has much weaker effects. Mechanistically, (R)-NNAL phosphorylates and deactivates LKB1 via the β-AR signaling in the LKB1 wild type NSCLC cell lines, contributing to the enhanced proliferation, migration, and drug resistance. Of note, NNK exposure also increases the phosphorylation of LKB1 in A/J mice. More importantly, human lung cancer tissues appear to have elevated LKB1 phosphorylation. Our results reveal, for the first time, that NNAL may promote lung cancer progression through LKB1 deactivation in an isomer-dependent manner.

## 1. Introduction

Lung cancer has been the leading cause of cancer-related deaths for decades [1–3]. It accounts for one in every five cancer deaths worldwide with about 160,000 deaths annually in the United States. While there are several other factors that may increase lung cancer risk, tobacco smoke is the main etiological factor associated with lung cancer development [4]. It is also associated with worse clinical outcome, including reduced therapeutic efficacy and shorter survival [5, 6]. For instance, the medium survival for non-smokers, former smokers, and active smokers among patients with non-small cell lung cancer (NSCLC) is 41.9, 22.6, and 14.7 months respectively [7]. However, 30 – 65% of NSCLC patients were active smokers at diagnosis and a significant portion of them continued to smoke (Table 1) [7–13]. It is therefore imperative to understand how tobacco use contributes to the worse clinical outcomes of lung cancer patients.

4-(Methylnitrosamino)-1-(3-pyridyl)-1-butanone (commonly known as “NNK”, Fig. 1A) is a tobacco specific lung carcinogen [14]. As a carcinogen, NNK is bioactivated by cytochrome P450 enzymes to induce DNA damage followed by subsequent mutations and carcinogenesis [15, 16]. NNK also promotes cell proliferation, enhances cell migration, and suppresses apoptosis in various cancer cell lines [17, 18]. Nicotinic acetylcholine receptors (nAChRs) [19] and β-adrenergic receptors (β-ARs) [20] have been suggested as the potential upstream targets of NNK. One key uncertainty of these in *vitro* results is their physiological relevance since NNK has a short-half life *in vivo* [21–24]. Indeed, NNK has never been detected in human biospecimens. Because of this, the metabolites of NNK have been used to investigate its human exposure and carcinogenic risk [25–28].

**Figure 1.**
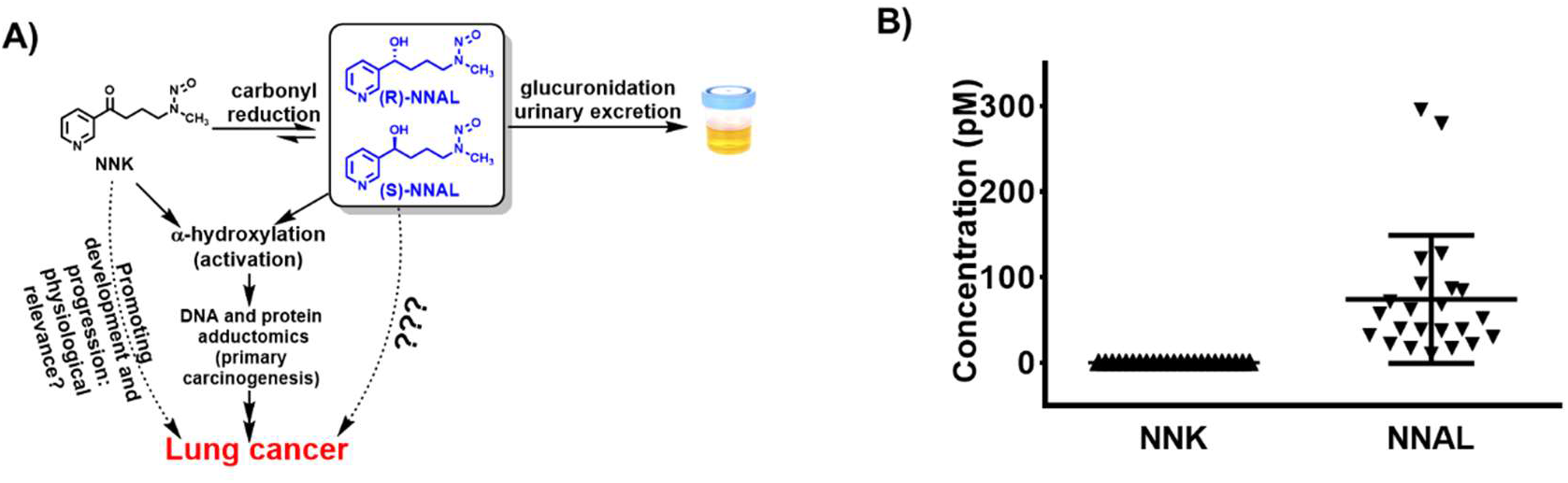
**A.** Simplified major pathways of NNK metabolism, carcinogenesis, and potential effects of NNK and NNAL on transformed lung cancer cells. **B.** Concentrations of NNK and NNAL in the plasma samples from human smokers.

**Table 1.**
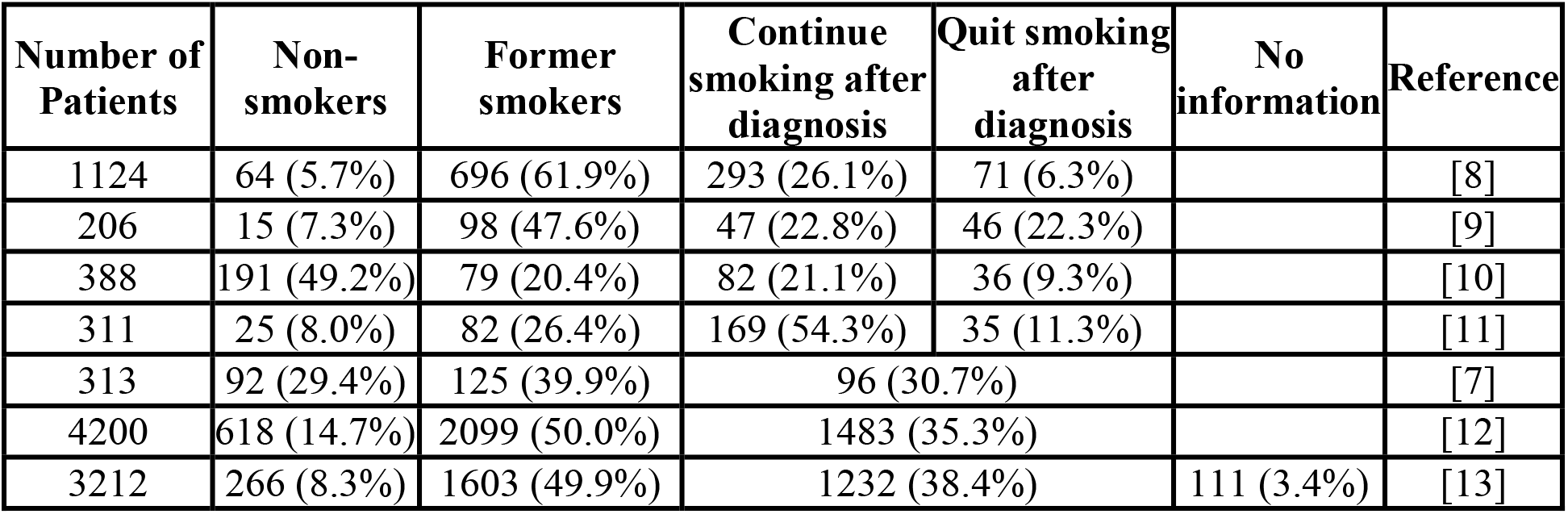
Smoking status among NSCLC patients.

| Number of Patients | Non-smokers | Former smokers | Continue smoking after diagnosis | Quit smoking after diagnosis | No information | Reference |
| --- | --- | --- | --- | --- | --- | --- |
| 1124 | 64 (5.7%) | 696 (61.9%) | 293 (26.1%) | 71 (6.3%) |  | [8] |
| 206 | 15 (7.3%) | 98 (47.6%) | 47 (22.8%) | 46 (22.3%) |  | [9] |
| 388 | 191 (49.2%) | 79 (20.4%) | 82 (21.1%) | 36 (9.3%) |  | [10] |
| 311 | 25 (8.0%) | 82 (26.4%) | 169 (54.3%) | 35 (11.3%) |  | [11] |
| 313 | 92 (29.4%) | 125 (39.9%) | 96 (30.7%) |  |  | [7] |
| 4200 | 618 (14.7%) | 2099 (50.0%) | 1483 (35.3%) |  |  | [12] |
| 3212 | 266 (8.3%) | 1603 (49.9%) | 1232 (38.4%) |  | 111 (3.4%) | [13] |

The major metabolite of NNK is 4-(methylnitrosamino)-1-(3-pyridyl)-1-butanol (commonly known as “NNAL”, Fig. 1A), which is formed via carbonyl reduction [29, 30]. Although structurally similar, NNAL has two key differences from NNK. First, NNAL has a much longer half-life *in vivo* relative to NNK [24]. In fact, NNAL is detectable in human urine samples weeks after the last tobacco exposure [31–33]. NNAL is also readily detected in blood samples from smokers [25]. For instance, Church *et al*. profiled serum levels of NNAL among 200 smokers selected from the Prostate, Lung, Colorectal, and Ovarian Cancer Screening Trial (PLCO). One hundred of these participants eventually developed lung cancer while the rest did not. Although the participants in these two groups were not rigorously matched in terms of age, gender and smoking history, NNAL concentrations were 92.4 ± 40.7 pM in the lung cancer cases and 77.4 ± 39.3 pM in the control groups. Second, NNAL has a chiral center (Fig. 1A). Its formation from NNK is catalyzed by a range of carbonyl reductases [26] and its elimination is mainly mediated through glucuronidation via UDP-glucuronosyltransferases (UGT) [27, 34]. Because of germline genetic variance in these metabolizing enzymes, smokers have a heterogeneous ratio of (R)-NNAL and (S)-NNAL [35]. These two enantiomers could have distinct biological activities. For instance, (S)-NNAL is much more carcinogenic than (R)-NNAL in A/J mice [34]. With the same dose treatment, (S)-NNAL resulted in much higher levels of DNA damage in the lung tissues and subsequently more lung adenoma formation than (R)-NNAL while (R)-NNAL was more efficiently eliminated via glucuronidation. These data argue that NNAL enantiomers may induce different biological effects and should be characterized as distinct individual entities. To date, the effect of NNAL on transformed lung cancer cells has not been reported. Such knowledge is important to help understand the reason for the worse outcome of lung cancer patients who continue to smoke.

In this study, we evaluated the effect of NNAL enantiomers on five human NSCLC cell lines at physiologically relevant concentrations. (R)-NNAL promoted cell proliferation, enhanced cell migration, and induced drug resistance while (S)-NNAL was substantially less effective. The effects of NNAL on cell migration and drug resistance required wild type liver kinase B1 (LKB1). Mechanistically, NNAL exposure, particularly the R enantiomer, led to LKB1 phosphorylation and deactivation through activating β-ARs. LKB1 phosphorylation was also observed in the lung tissues of A/J mice upon NNK exposure. Human lung cancer tissues had substantially higher levels of phosphorylated LKB1 relative to the paired normal lung tissues. In summary, NNAL, particularly (R)-NNAL, deactivates LKB1 through β-ARs in NSCLC cancer cell lines. Such LKB1 deactivation confers drug resistance and promotes invasion. In addition, LKB1 loss-of-function human lung cancers may be highly prevalent via phosphorylation due to common tobacco exposure in addition to the mutational deactivation. Overall, our results depict a novel mechanism through which active smoking may contribute to the worse outcome of lung cancer patients.

## 2. Results

### 2.1. Detection and quantification of NNK and NNAL in human blood samples from smokers and non-smokers

To determine the physiologically relevant compound(s) and concentrations for our studies, we quantified NNK and NNAL in the plasma from smokers (n = 46) and non-smokers (n = 3) via an established liquid chromatography with tandem mass spectrometry (LC-MS/MS) method [36]. The smoking status of the plasma donors was confirmed by measuring their urinary total nicotine equivalents (TNE). Some of the results have been published [37, 38]. Consistent with its short half-life, NNK was not detected in any of these samples (Fig. 1B). NNAL was not detectable in the plasma samples from non-smokers (not shown) while it was readily detected in the plasma samples from smokers (Fig. 1B). The plasma concentration of NNAL ranged between 10.4 pM to 296.0 pM with a mean value of 59.7 ± 61.1 pM. NNAL was therefore evaluated in our *in vitro* study instead of NNK.

### 2.2. The effects of NNAL enantiomers on cell proliferation, migration and drug resistance in NSCLC cancer cells with different LKB1 status

The concentrations of NNAL detected in the plasma samples from smokers were in the picomolar range, consistent with those reported in the literature [25, 39, 40]. NNAL concentrations in the lung are expected to be much higher because of the direct exposure of lung to tobacco smoke. We proposed that NNAL concentrations between 1-100 nM are physiologically relevant, and this range was therefore used in our subsequent *in vitro* studies.

NNAL exposure in H1299 and A549 cells resulted in no detectable (*O*^6^-mG (data not shown), suggesting the lack of NNAL bioactivation and associated carcinogenesis. However, NNAL at 10 nM significantly increased cell proliferation (Fig. 2A). When the two enantiomers of NNAL were evaluated, (R)-NNAL recapitulated this activity in both cell lines while (S)-NNAL had minimal effects (Fig. 2B). Similarly, (R)-NNAL significantly increased colony formation in H1299 and A549 cells while (S)-NNAL was not effective (Fig. 2C). The potential of NNAL enantiomers on cell migration was evaluated via wound healing assay (Fig. 2D) and trans-well assays (Fig. 2E) in H1299 and A549 cells. In both assays, (R)-NNAL substantially increased cell migration while (S)-NNAL had little effects in H1299 cells. Intriguingly, NNAL treatment had no effects in A549 cells (Fig. 2D and 2E). Lastly, the effect of NNAL on the cytotoxicity of gemcitabine and cisplatin was evaluated via a cell viability assay. (R)-NNAL significantly reduced the cytotoxicity of gemcitabine and cisplatin. (S)-NNAL conferred less resistance compared with (R)-NNAL (Fig. 2F). Both NNAL enantiomers failed to induce resistance in A549 cells (Fig. 2F).

**Figure 2.**
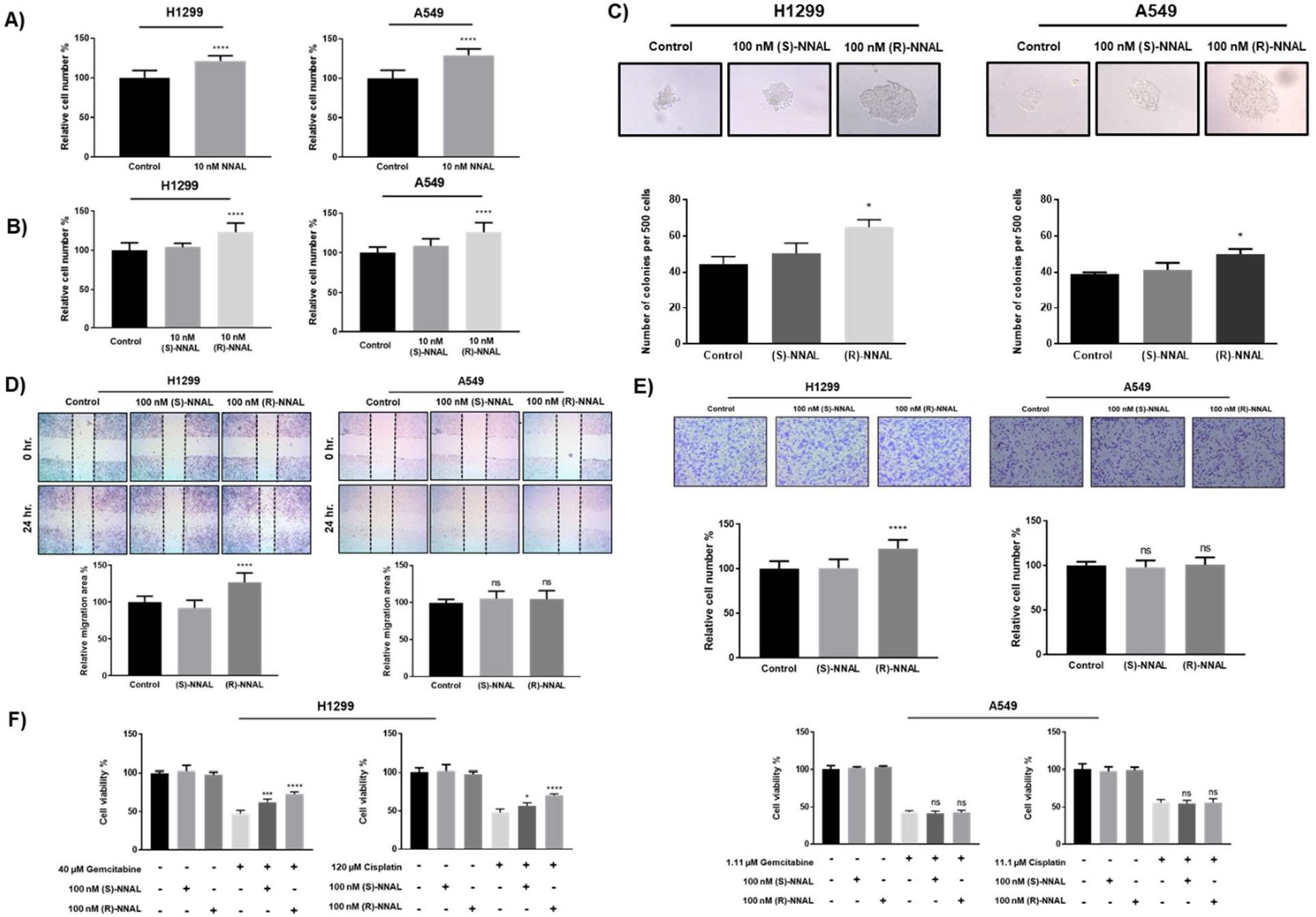
Effect of NNAL enantiomers on malignant phenotypes in H1299 and A549 lung cancer cells. **A.** Effect of NNAL (10 nM) on cell proliferation. **B.** Effect of NNAL enantiomer (10 nM) on cell proliferation. **C.** Effect of NNAL enantiomers (100 nM) on colony formation. **D.** Effect of NNAL enantiomers (100 nM) on cell migration via wound healing assay. **E.** Effect of NNAL enantiomers (100 nM) on cell migration via transwell assay. **F.** Effect of NNAL enantiomers (100 nM) on conferring drug resistance to gemcitabine (40 μM) or cisplatin (120 μM) treatment. *, P<0.05; **, P<0.01; ***, P<0.001.

Although there are many molecular and genetic differences between H1299 and A549 cell lines that could account for the observed differences in cellular migration and drug resistance, we focused on LKB1 because of its importance in lung cancer development [41–48] and its different status in H1299 (wild-type) and A459 (mutational deactivation). Moreover, LKB1 is a potential down-stream target for β-ARs, which NNK has been reported to activate [19, 20]. We therefore evaluated the effect of (R)-NNAL in HCC827 (*LKB1* WT), H1975 (*LKB1* WT) and H460 (*LKB1* mutant) cells. As have been observed in H1299 and A549, (R)-NNAL enhanced cell proliferation in all of these cell lines (Fig. S1). While (R)-NNAL reduced the sensitivity of HCC827 and H1975 cells to cisplatin and gemcitabine, it failed to reduce the sensitivity of H460 to these therapies (Fig. S2). Together, these data suggest that the status of *LKB1* in NSCLC cancer cell lines may be critical to the detrimental effects of NNAL, particularly in cell migration and drug resistance.

### 2.3. The role of LKB1 on NNAL-mediated cell migration and drug resistance in NSCLC cells

To characterize the role of LKB1 in mediating the cellular effects of NNAL, an *LKB1*-knockout H1299 cell line was generated using a CRISPR knockout approach (LKB1-KO H1299, Fig. 3A). As expected, (R)-NNAL failed to facilitate cell migration in LKB1-KO H1299 cells (Fig. 3B) and did not induce resistance to gemcitabine nor cisplatin (Fig. 3C). Similarly, WT LKB1 was knocked into A549 cells (Fig. 3D). (R)-NNAL was able to promote cell migration (Fig. 3E) and confer drug resistance in A549 cells with LKB1 knock-in (Fig. 3F). Altogether, these data suggest that LKB1 plays a key role in NNAL-mediated migration and drug resistance in *LKB1* WT lung cancer cells and the loss of LKB1 significantly reduced the impact of (R)-NNAL exposure.

**Figure 3.**
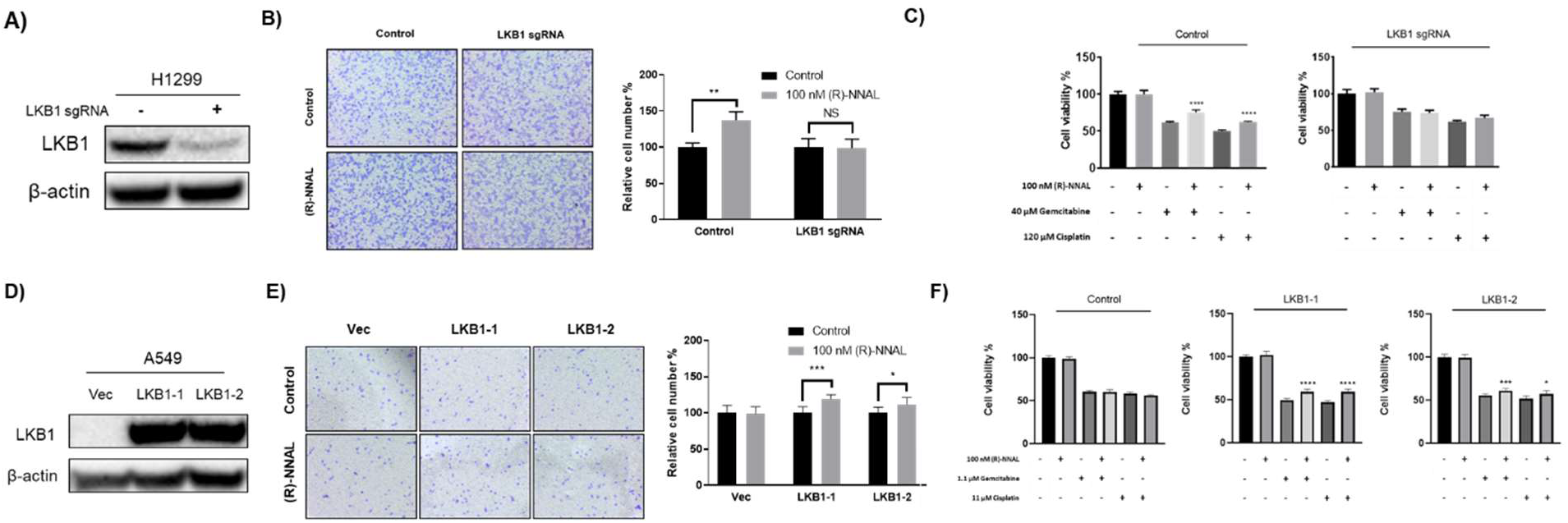
The function of LKB1 on malignant phenotypes promoted by NNAL exposure. **A.** Expression of LKB1 in H1299 LKB1 knockout cells. **B.** Effect of (R)-NNAL on cell migration in H1299 LKB1 knockout cells. **C.** Effect of (R)-NNAL on the cytotoxicity of gemcitabine and cisplatin in H1299 LKB1 knockout cells. **D.** Expression of LKB1 in A549 LKB1-knockin cells. **E.** Effect of (R)-NNAL on cell migration in A549 LKB1-knockin cells. **F.** Effect of (R)-NNAL on the cytotoxicity of gemcitabine and cisplatin in A549 LKB1-knockin cells. *, P<0.05; **, P<0.01; ***, P<0.001.

### 2.4. The effect of NNAL enantiomers on LKB1 phosphorylation in lung cancer cells

We next characterized the effect of NNAL enantiomers on LKB1 phosphorylation in H1299, H1975, and HCC827 cells. Increased deactivating phosphorylation of LKB1 at Ser428 was observed in all of these cells with greater increase upon the (R)-NNAL exposure than the (S)-enantiomer (Fig. 4A). The time course of LKB1 phosphorylation by (R)-NNAL was characterized in H1299 cells (Fig. 4B). (R)-NNAL treatment also led to a significant reduction in phosphorylated AMPK, and increase in phosphorylated mTOR and 4E-B1 (Fig. 4C). Since AMPK, mTOR and 4EB-P1 are downstream proteins of LKB1, the reduction in AMPK phosphorylation and increase in mTOR and 4E-BP1 phosphorylation suggest the deactivation of LKB1, consistent with its increased phosphorylation. (R)-NNAL treatment also reduced cleaved PARP caused by cisplatin treatment, had little effect on cisplatin-induced DNA damage, and may slightly reduce the level of Bim protein (Fig. 4D and Fig. 4E), which may explain the reduced sensitivity of H1299 cells to cisplatin treatment in the presence of (R)-NNAL. In addition, (R)-NNAL treatment resulted in a slight increase in the level of PCNA (Fig. 4F), potentially accounting for its stimulation of cell proliferation.

**Figure 4.**
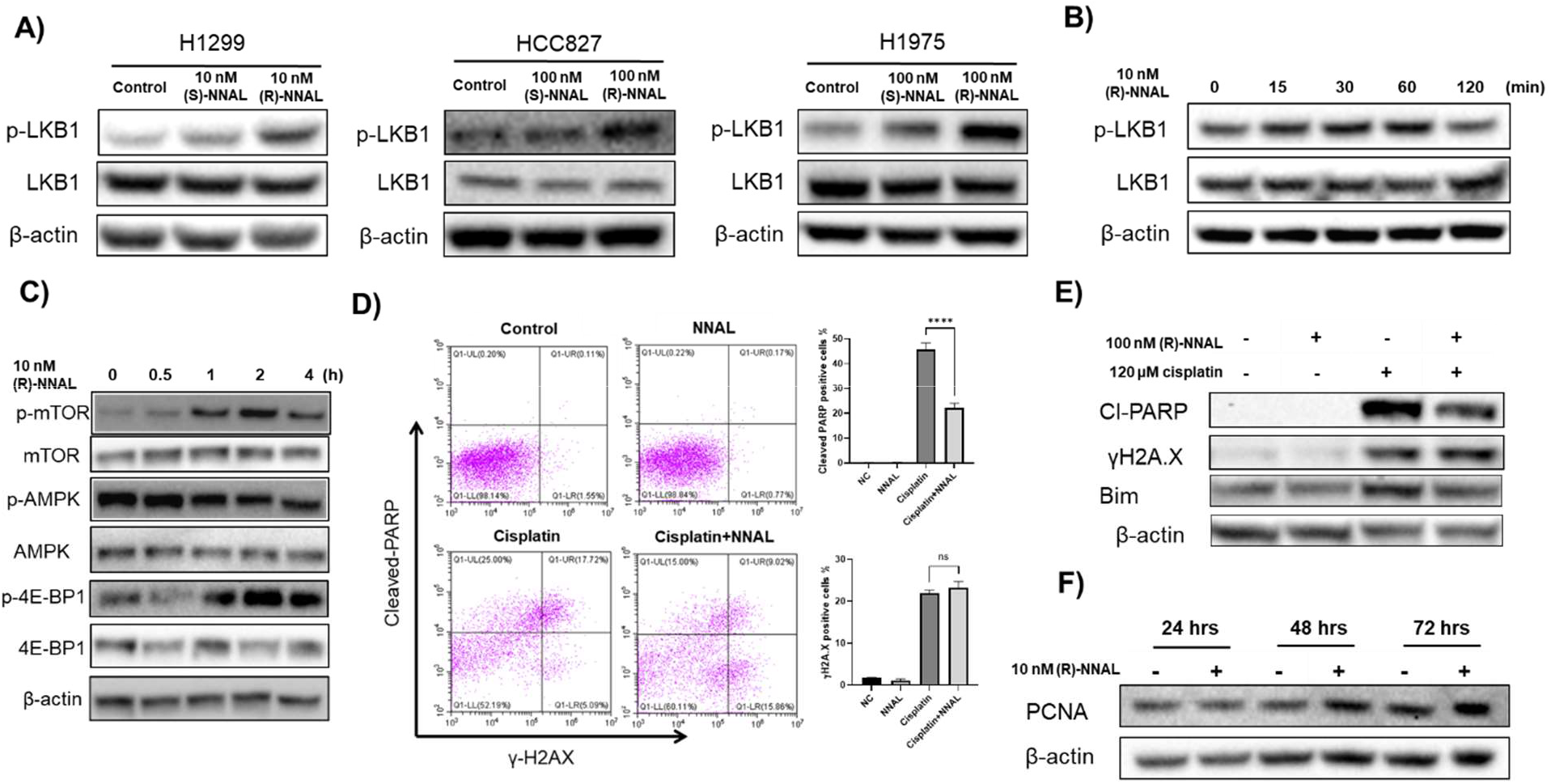
Effects of NNAL on LKB1 phosphorylation and associated signaling in *LKB1* WT lung cancer cells. **A.** The effects of (R)- and (S)-NNAL on LKB1 deactivating phosphorylation at Ser428. Cells were treated with 10 nM NNAL for 30 min. **B.** Time course effect of (R)-NNAL on the phosphorylation of LKB1 at Ser428 (H1299). **C.** Time course effect of (R)-NNAL on AMPK, mTOR and 4EBP1 phosphorylation in H1299. **D.** and **E.** Effect of (R)-NNAL on cisplatin induced DNA damage and PARP cleavage. **F.** Time course effect of (R)-NNAL exposure on PCNA levels in H1299.

### 2.5. The upstream signaling responsible for NNAL-mediated LKB1 phosphorylation

Upon establishing the role of LKB1, we explored the potential upstream targets of NNAL responsible for LKB1 phosphorylation and associated phenotypes. NNK has been reported to act as an agonist for nAChRs [19] and β-ARs [20]. This could result in the activation of protein kinase A (PKA) via intracellular calcium influx and cAMP synthesis, which would phosphorylate and deactivate LKB1 [49]. First, we found (R)-NNAL could promote PKA-Cα nucleus translocation in H1299 (Fig. 5A). And then, we utilized a panel of pharmacological inhibitors to probe the relevance of these potential upstream signaling processes, including propranolol (a β-AR antagonist), nifedipine (a calcium channel blocker), H89 (a PKA inhibitor) and yohimbine (an α2-AR antagonist as a control). We evaluated their effects on NNAL-induced proliferation in H1299 cells (Fig. 5B). At non-cytotoxic concentrations, each pharmacological inhibitor, except yohimbine, effectively blocked (R)-NNAL-induced enhanced proliferation. Similarly, these pharmacological inhibitors, with the exception of yohimbine, effectively blocked (R)-NNAL induced resistance against gemcitabine or cisplatin in H1299 cells (Fig. 5C). Consistently, each pharmacological inhibitor, with the exception of yohimbine, reduced the phosphorylation of LKB1 (Ser428) induced by (R)-NNAL exposure (Fig. 5B). Overall, these data delineate the signaling process of LKB1 deactivation by NNAL.

**Figure 5.**
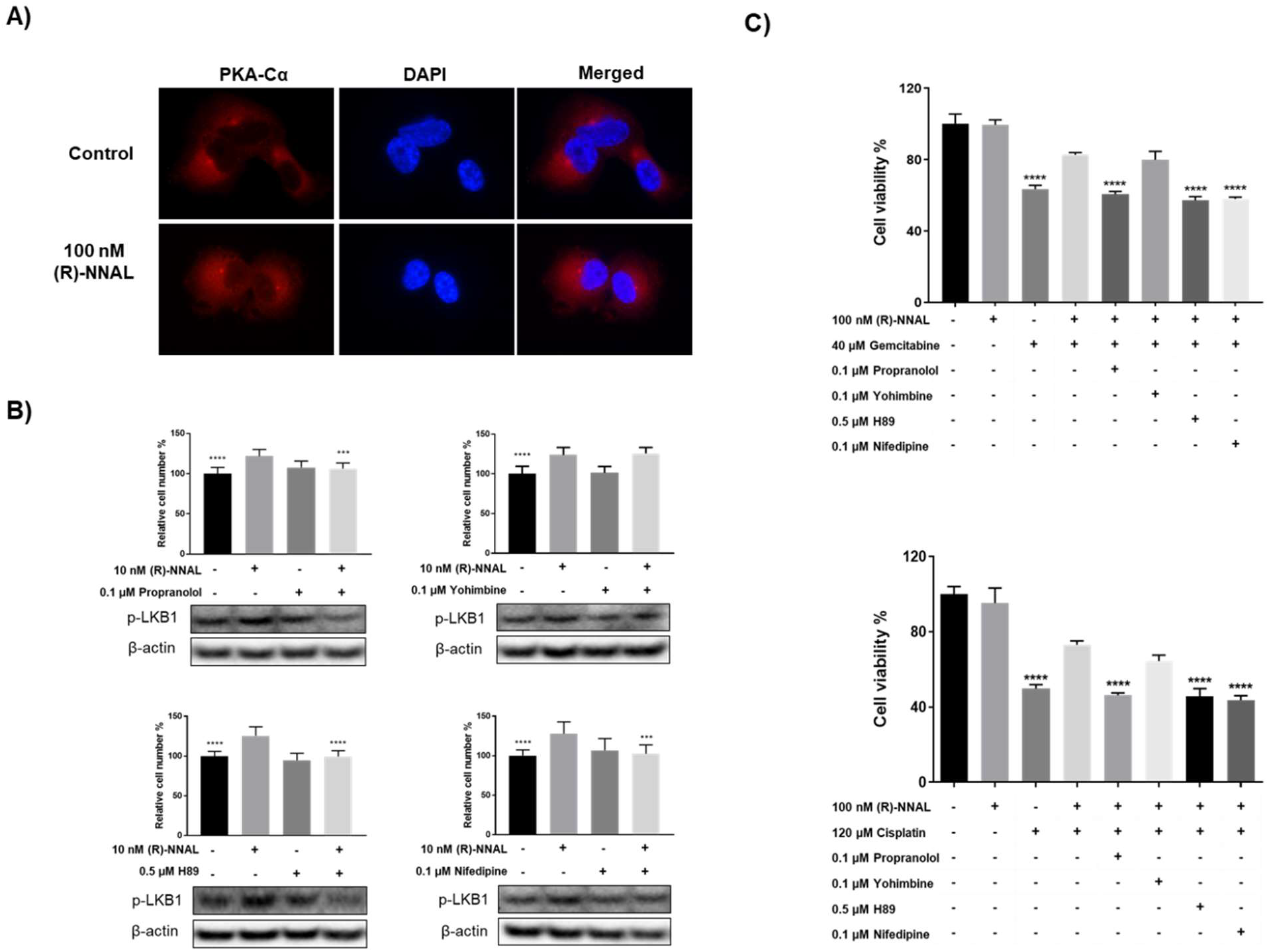
The potential upstream signaling events governing NNAL-promoted phenotypes with pharmacological inhibitors. **A.** Effect of (R)-NNAL on PKA-Cα nucleus translocation in H1299. Cells were treated with 100 nM (R)-NNAL for 60 min. DAPI was used to stain the nucleus. **B.** Effect of inhibition of β-ARs (propranolol), Ca^2+^ channels (nifedipine), PKA (H89) and α-ARs (yohimbine) on NNAL-promoted cell proliferation and LKB1 phosphorylation (Ser428). H1299 cells were co-treated with 10 nM (R)-NNAL and 0.1 μM nifedipine, 0.1 μM propranolol, 2.5 μM bupropion, 0.5 μM H89 or 0.1 μM yohimbine for 6 days. **C.** Effect of inhibition of β-ARs (propranolol), Ca^2+^ channels (nifedipine), PKA (H89) an α-ARs (yohimbine) on NNAL-promoted resistance to gemcitabine and cisplatin. H1299 cells were co-treated with 40 μM gemcitabine or 120 μM cisplatin along inhibitors. *, P<0.05; **, P<0.01; ***, P<0.001.

### 2.6. The effect of prolonged (R)-NNAL exposure on H1299 cells

The lung tissue of smokers may be chronically exposed to NNAL due to the habitual use of tobacco and the slow elimination of NNAL. We therefore evaluated the effect of long-term (R)-NNAL exposure on H1299 cells. Specifically, H1299 cells were cultured with (R)-NNAL (1 nM) for 60 days. Then, in the absence of NNAL, the phosphorylation status of LKB1 was characterized and cell proliferation, colony formation, cell migration and the sensitivity of such cells to gemcitabine and cisplatin treatment was evaluated. Long-term NNAL exposure resulted in a substantial increase in LKB1 (Ser428) phosphorylation even in the absence of NNAL (Fig. 6A). These H1299 cells proliferated faster, supported by the cell proliferation data (Fig. 6B) and colony formation data (Fig. 6C). And these H1299 cells were also significantly less sensitive to cisplatin and gemcitabine treatment in the absence of NNAL (Fig. 6D). These data suggest that the effect of long-term NNAL exposure on LKB1 deactivation and drug resistance could be long-lasting. Interestingly, the addition of NNAL to such cells failed to further enhance drug resistance (data not shown). Mechanically 60 days exposure to 1 nM (R)-NNAL has little effect on cisplatin induced DNA damage indicated by the level of γH2A.X, and significantly reduced PARP cleavage (Fig. 6E.). In addition, H1299 cell migration was also enhanced upon 60 days exposure to (R)-NNAL (Fig. 6F. and Fig. 6G).

**Figure 6.**
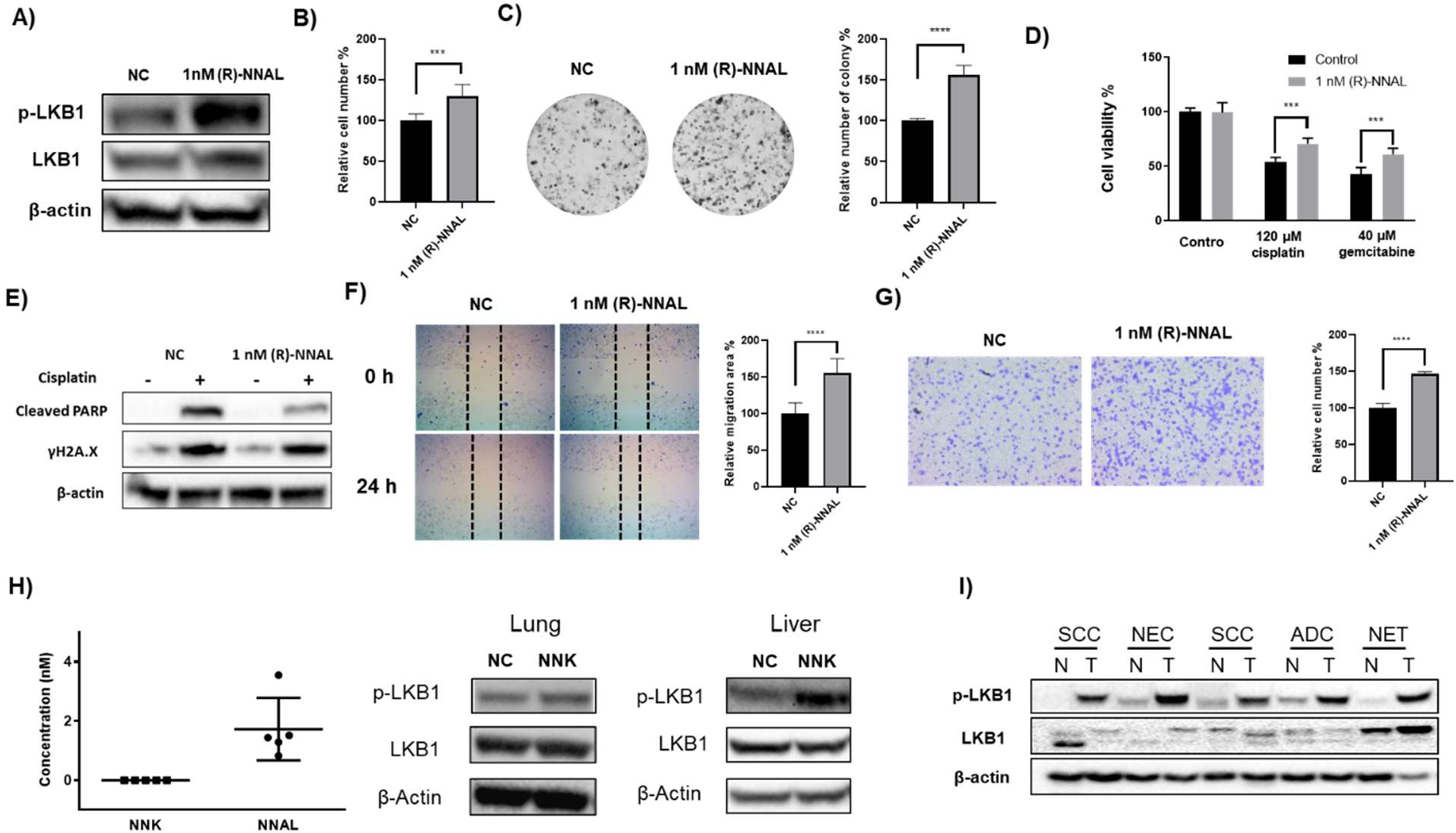
LKB1 phosphorylation status *in vitro, in vivo* and in clinical samples with potential chronic NNAL exposure. Effect of 60-day (R)-NNAL exposure on cell proliferation (**A**), colony formation (**B**), sensitivity to gemcitabine and cisplatin treatment (**C**), cell migration (**D** and **E**), and LKB1 phosphorylation (**F**) in H1299 cells. H1299 cells was treated with (R)-NNAL (1 nM) for 60 days and no additional (R)-NNAL was added when running these assay. **G.** Concentrations of NNK and NNAL in mouse serum (n = 5) and LKB1 status in the lung and liver tissues of A/J mice upon 4-week NNK exposure. A/J mice were given NNK in drinking water (40 ppm) for 4 weeks. **H.** Status of LKB1 in normal (N) or tumor (T) tissues of five lung cancer patients (SCC: squamous cell carcinoma; ADC: adenocarcinoma; NEC: neuroendocrine carcinoma; NET: neuroendocrine carcinoid).

### 2.7. The effect of NNK exposure in A/J mice on LKB1 phosphorylation

To explore whether NNK induces LKB1 phosphorylation in vivo, a pilot study in A/J mice was performed. In this study, NNK was administered in drinking water at a concentration of 40 ppm. This treatment regimen mimics the chronic exposure of NNK among smokers although liver instead of lung is the tissue of main exposure. The dose of NNK (40 ppm) in mice is comparable to the level of NNK exposure among heavy smokers [50]. A similar treatment regimen has been demonstrated to induce lung adenoma formation in A/J mice [51–53]. The lung and liver tissues were collected after a 4-week NNK exposure. Again, NNK was not detectable in any serum samples while NNAL was detected all (Fig. 6H), consistent with human data. The serum concentration of NNAL in the mice ranged between 0.83 – 3.55 nM, similar to the concentration used in our *in vitro* studies. NNK treatment substantially increased LKB1 phosphorylation in the liver tissues with a slight increase in the lung tissues (Fig. 6H), indicating the deactivation of LKB1 in A/J mice upon NNK exposure. The higher levels of LKB1 phosphorylation in the liver tissues than the lung tissues in this model may be caused by the NNK drinking water regimen that the liver tissues have higher exposure to NNK than the lung tissues. In human smokers, the lung tissues have higher exposure to NNK that may favor LKB1 phosphorylation in the lung tissues.

### 2.8. LKB1 status in lung cancer tissues

To explore the potential clinical significance of LKB1 phosphorylation by NNAL, we characterized the phosphorylation status of LKB1 protein in five lung cancer tissues in comparison to the normal tissues from the same patients (Fig. 6I). Although there are variations and no obvious patterns in the total protein levels of LKB1 between the normal and cancer tissues, p-LKB1 (Ser428) were substantially higher in the cancer tissues relative to the normal tissues irrespective of the lung cancer pathology.

## 3. Discussion

Clinical management of lung cancer has not been very successful and the overall survival from lung cancer remains frustratingly low [1–3]. There are many contributing factors, including late diagnosis and higher risk of drug resistance and metastasis [54–56]. At the same time, many lung cancer patients are active smokers at the time of diagnosis and a significant portion of them continue to smoke, which is associated with worse outcomes [5, 6]. NNK has been proposed as a contributing factor because it could enhance lung cancer proliferation and survival, and promote metastasis *in vitro* and *in vivo* [17, 18, 57, 58]. Potential mechanisms have been characterized *in vitro*, including the activation of CREB, ERK1/2, and Akt with nAChRs and β-ARs as the upstream targets [19, 20, 58, 59]. These mechanistic studies, however, may have limited physiological relevance because NNK is not detectable in human plasma samples. Its major metabolite, NNAL, on the other hand, is detected in the plasma samples from all smokers in our study with a concentration approaching 300 pM. Given that lung tissue has the highest exposure to tobacco smoke, NNAL between 1 and 100 nM *in vitro* is likely physiologically relevant. Within this concentration range, NNAL enhanced cell proliferation in all five NSCLC cancer cell lines tested, with (R)-NNAL being more potent than (S)-NNAL. NNAL also promoted cell migration and induced drug resistance in NSCLC cell lines that have wild-type LKB1. Such effects were also more pronounced with (R)-NNAL than (S)-NNAL. These results suggest that the detrimental effects of NNAL may vary among smokers because of genetic polymorphisms in NNAL metabolizing enzymes, such as carbonyl reductases and UGTs. It should also be noted that although the stimulating effects by NNAL on proliferation, migration and resistance are not very strong under our experimental conditions, the cumulative impact should not be underestimated given the chronic exposure of the lungs to NNAL among smokers. Indeed, a 60-day exposure of H1299 cells to (R)-NNAL (1 nM) resulted in significant enhancing of cell proliferation, migration and drug resistance in combination with LKB1 phosphorylation even in the absence of NNAL.

LKB1 mutational deactivation has been observed in 10 – 30% of lung cancer patients of different pathological subtypes [41–44]. Results from a number of genetic mouse models strongly indicate that LKB1 inactivation plays an important role in lung cancer initiation, development, and progression [46–48, 60]. Indeed, lower levels of LKB1 expression has been reported to be associated with higher recurrence in NSCLC [61] and loss of LKB1 has been discussed beyond just mutations [62]. Upon analyzing a limited number of lung cancer tissues, we observed enhanced LKB1 phosphorylation in lung cancer tissues compared with paired normal lung tissues. These data suggest that the function of wild-type LKB1 protein in lung cancers may be compromised and the potential contribution of LKB1 deactivation to human lung cancer could be substantially higher than its mutational frequency, something that warrants future investigation. It is therefore of great importance to understand how wild-type LKB1 is phosphorylated in lung cancer patients. Our results showed for the first time that NNAL, particularly (R)-NNAL, induces LKB1 phosphorylation (Ser428) in NSCLC cancer cells. Since a 60-day NNAL exposure resulted in phosphorylated LKB1 even upon NNAL removal, prior or active tobacco use among former and current smokers, respectively, could contribute to the phosphorylation and deactivation of LKB1 in human lung cancer tissues. This was further supported by our A/J mouse data that a 4-week NNK exposure resulted in LKB1 phosphorylation in the lung tissues. Based on our results with pharmacological inhibitors, β-ARs are the potential up-stream target(s) for NNAL that then activate PKA, leading to LKB1 phosphorylation (Fig. 7). Other agonists for β-ARs, such as mental stress-related stress hormones norepinephrine and epinephrine, may also deactivate LKB1 in humans, which again warrants future investigation. Indeed, nicotine exposure and mental stress have also been documented as potential factors contributing to the worse outcome of lung cancer patients [63, 64].

**Figure 7.**
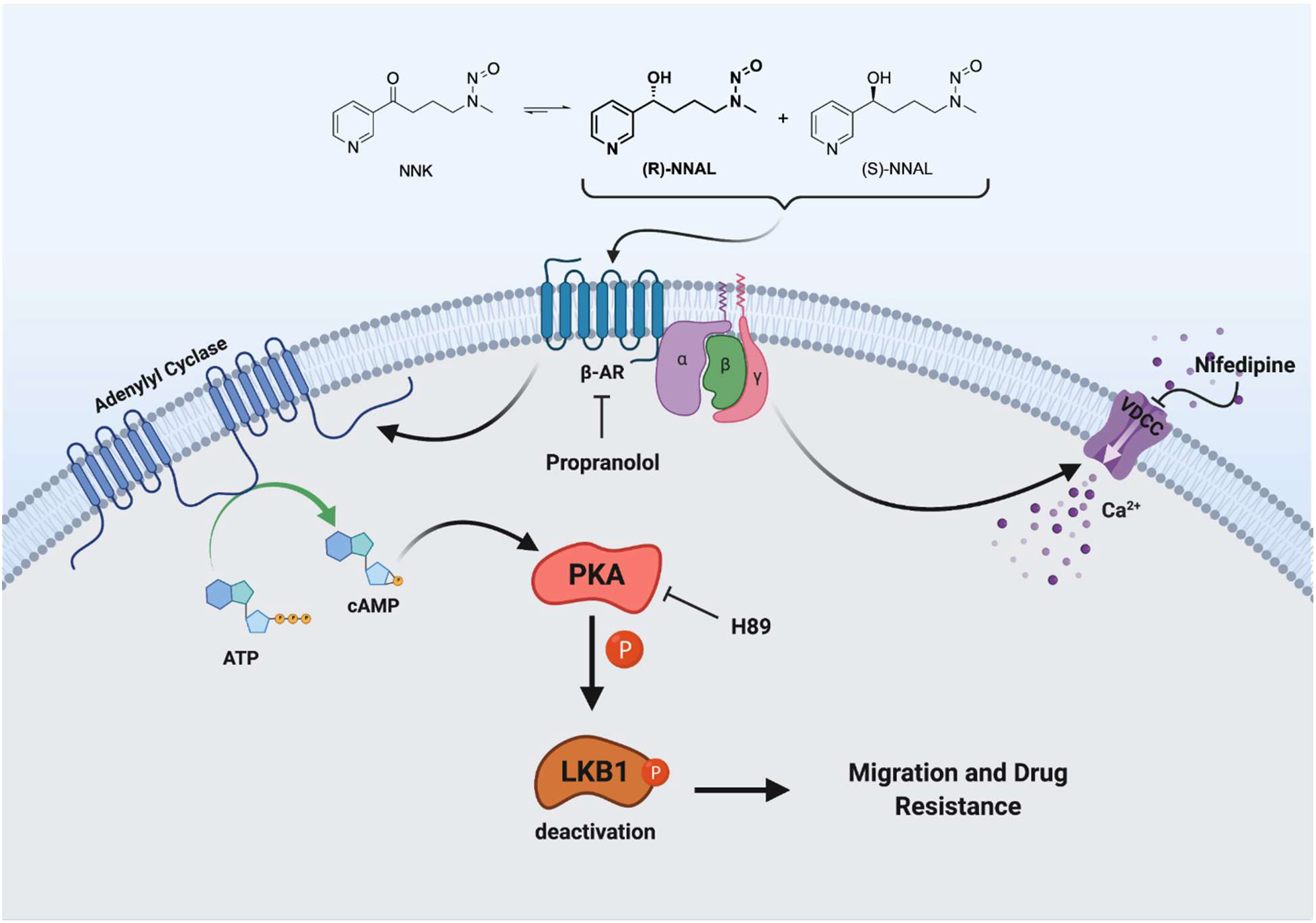
Proposed mechanisms of NNAL in promoting progression of lung cancer cells with wild type LKB1.

Of note, (R)-NNAL stimulated cell proliferation in all NSCLC cancer cells irrespective of LKB1 status, suggesting that NNAL modulates signaling mechanisms independent of LKB1. NNK has been reported to activate CREB, a master oncoprotein [57, 65, 66], to promote progression in established tumors. CREB activation is dominantly mediated via PKA as well. We therefore evaluated the effect of NNAL on CREB phosphorylation in all five NSCLC cells. NNAL rapidly activated CREB in these cell lines independent of LKB1 status with (R)-NNAL being more potent than (S)-NNAL (Fig. S3). Thus, increased CREB phosphorylation and activation may contribute to the increased proliferation of NSCLC cancer cells induced by NNAL.

In summary, our results show that NNAL can deactivate LKB1 in lung cancer cells at physiologically relevant concentrations in an isomeric dependent manner. Such deactivation may be of great clinical relevance given the tumor suppressive functions of LKB1 in lung cancer initiation, development and progression and the high prevalence of tobacco exposure among lung cancer patients. Further in vivo and clinical studies are warranted to validate NNAL’s tumor promoting effects, its contribution to LKB1 deactivation and the worse clinical outcome of lung cancer patients who continue to smoke.

## 4. Materials and Methods

**Caution:** Both NNK and NNAL are highly carcinogenic. They should be handled in a well-ventilated hood with extreme care, and with proper personal protective equipment.

### 4.1. Chemicals and Reagents

NNK, [^13^C_6_]NNK, [^13^C_6_]NNAL, [CD_3_]*O*^6^-mG were purchased from Toronto Research Chemicals (Toronto, Ontario, Canada). (±)-Propranolol hydrochloride, bupropion and H89 were purchased from Sigma-Aldrich (St. Louis, MO, USA). Nifedipine was purchased from Alfa Aesar (Ward Hill, MA, USA). Yohimbine hydrochloride was purchased from Acros Organics (Morris, NJ, USA). All reagents were used without further purification.

### 4.2. Human samples

Plasma samples from active smokers were collected from a clinical trial previously conducted at the University of Minnesota [36] and from active smoking population controls in the NCI-MD Lung cancer Case Control Study [67]. Plasma samples from non-smokers were purchased from Bioreclamation IVT (Baltimore, MD). Demographic information of these donors has been published before [37, 68]. Paired normal and cancerous lung tissues from five patients were acquired from University of Florida CTSI Biorepository. The protocols for human sample use were reviewed and approved by Institutional Review Boards (IRB) at the University of Florida.

### 4.3. NNAL synthesis, chiral resolution, and characterization

Racemic NNAL was synthesized from NNK via sodium borohydride reduction. (R)- and (S)-NNAL were separated from the racemic mixture via chiral chromatography by a contract service from Kermanda Biotech Co Ltd. (Shanghai, China). Racemic, (R)- and (S)-NNAL were characterized by ^1^H-NMR and HPLC with > 95% purity. The chirality of the two enantiomers, (R)- and (S)-NNAL, was assigned on the basis of reported optical rotations of NNAL [69].

### 4.4. NNK and NNAL quantification in human plasma samples and mouse serum samples

The concentrations of NNK and NNAL in human plasma and mouse serum samples were quantified following a previously reported mass spectrometry method [70].

### 4.5. Cell lines and culturing conditions

H1299, H1975, HCC827, H460 and A549 cells were purchased from ATCC (Manassas, VA). H1299, A549 and H460 were authenticated via the Cell Line Authentication Service provided by Genetica DNA Laboratories (Burlington, NC). H1975 and HCC827 were authenticated by ATCC. All of these cell lines were confirmed to be free from mycoplasma infection. H1299, H1975, HCC827 and H460 cells were maintained in RPMI 1640 medium supplemented with 10% FBS (Gibco). A549 cells were maintained in DMEM medium supplemented with 10% FBS. All cells were cultured in a 37 °C, 5% CO_2_ atmosphere. H1299 LKB1 knockout was reported before [71]. For A549 LKB1 knock-in, STK11(LKB1) gene was subcloned into PLX-304 vector. Lentivirus production was performed using psPAX2 (Addgene#12260) and pMD2.G (Addgene#12259) as previously described [72]. Single clones of cells expressing LKB1 were selected using blasticidin (5ug/ml) and LKB1 expression was confirmed by western blot.

### 4.6. Analysis of *O*^6^-methylguanine (*O*^6^-mG) in H1299 cells upon NNAL treatment

Among the various forms of DNA damages caused by NNK and NNAL, (*O*^6^-mG was the most abundant in A/J mouse lung tissues [73] and of comparable abundance to other types of DNA damage in F344 rat lungs [74] although such DNA damage has not been detected in human lung tissues. We therefore quantified *O*^6^-mG in H1299 cells upon NNAL exposure (100 nM) using an established mass spectrometry method [75].

### 4.7. Cell counting assay

Cell proliferation was determined using a cell count assay. Briefly, 5,000 cells/well were seeded in a 24-well plate with 10% FBS medium. After overnight incubation, medium was replaced with 0.5% FBS medium containing NNAL. After a 6-day incubation, cells were trypsinized and cell numbers were determined using the Bio-Rad Automated Cell Counter.

### 4.8. Colony formation assay

Cell proliferation was also determined using colony formation assay. Briefly, cells were plated in a 24-well plate (500 cells/well) in 0.5% FBS medium with or without NNAL. The number of colonies was counted after a 7-day incubation under the microscope.

### 4.9. Wound healing assay

Cell migration was measured using the wound healing assay. Briefly, cells were seeded into a six-well plate and allowed to grow to ~90% confluency. After starvation with FBS-free medium for 48 h, cell monolayers were wounded with a 200-μL pipette tip. Wounded monolayers were washed three times with PBS and incubated in serum-free medium with different concentrations of NNAL for 24 h. Cells were monitored under a microscope equipped with a camera. The wound area was quantified using Image J software.

### 4.10. Transwell assay

Cell migration was also evaluated with the transwell migration assay using 6.5 mm diameter inserts (Corning) with 8 μm pore size. The inserts were plated in a 24-well plate with 600 μL 10% FBS medium. Briefly, 30,000 cells in 200 μL serum free medium with 100 nM (R)- or (S)-NNAL were seeded into each insert. After incubation at 37 °C for 24 h, the cells in the upper surface of the membrane were removed with a cotton swab. Cells in the lower chamber were fixed with 70% ethanol and stained with 0.2% crystal violet (Sigma-Aldrich in St. Lewis, MO, USA). Images were taken with an inverted microscope and the number of cells was quantified using ImageJ.

### 4.11. Cell viability assay

Drug resistance was evaluated via a cell viability assay. Briefly cells were plated in 96-well plates (5,000 cells/well) with 10% FBS medium. After attachment, cells were treated with the test compounds at the specified concentrations or combinations in triplicate with 0.5% FBS medium. The relative cell viability in each well was determined after 72 h treatment using the MTT assay (Life Technologies).

### 4.12. NNK exposure in A/J mice

Female A/J mice (5-6 weeks of age) were purchased from the Jackson Laboratory (Bar Harbor, ME) and maintained in specific pathogen-free facilities, according to animal welfare protocols approved by Institutional Animal Care and Use Committee at the University of Florida. After 1-week acclimation, mice were weighed, randomized into two groups (n = 5) and switched to AIN-93G powdered diet, defined as Day 1. From Day 1, mice in the control group were given regular drinking water while the NNK group was given NNK in drinking water (40 ppm). Mice were euthanized 4 weeks after NNK exposure. The lung and liver tissues were harvested, snap-frozen in liquid N_2_ and stored at −80 °C until protein analysis. Serum was collected for NNK and NNAL detection.

### 4.13. Western blotting

Whole cell lysates from H1299, H1975, HCC827, A549 and H460 cells were prepared in RIPA lysis buffer. Protein lysates from human and mouse lung tissues were prepared similarly. Briefly, 20 mg tumor or normal tissue was homogenized in 250 μL RIPA buffer and the supernatant was collected after centrifugation at 13,000 g for 15 min at 4 °C. The concentration of protein in each sample was quantified using BCA assay. Forty - sixty μg of protein from each sample was denatured in SDS-PAGE sample buffer and resolved on 4-12% Bis-Tris PAGE gels. The separated proteins were transferred to Polyvinylidene difluoride (PVDF) membrane followed by blocking with 5% non-fat milk powder (w/v) in Tris-buffered saline (10 mM Tris–HCl, pH 7.5, 100 mM NaCl, 0.1% Tween-20) for 1 h at room temperature. After blocking, the membranes were probed with desired primary antibodies overnight at 4 °C followed by appropriate peroxidase-conjugated secondary antibody for 2 h at room temperature and visualized by the Bio-Rad ChemiDoc Imaging system. To ensure equal protein loading, each membrane was stripped with Restore Western Blot stripping buffer (Thermo Scientific) and re-probed with β-actin antibody. Detailed information on antibodies is in Table S1.

### 4.14. Immunofluorescence staining

Treated cells were fixed with 4% paraformaldehyde for 15 min and permeabilized with 1% Triton X-100 in PBS for 10 min, followed by blocking with 5% BSA in PBS for 1 h. After blocking, cells were incubated with PKA-Cα antibody overnight at 4 °C and secondary antibody for 1 h at room temperature. Nucleus were stained with Dapi for 15 min. Cells were imaged using Fluorescence microscopy (Nikon Ti2, Japan).

### 4.15. Flow cytometry

Detection of γH2A.X and cleaved PARP protein level in cisplatin treated H1299 with/without (R)-NNAL were performed using Apoptosis, DNA Damage and Cell Proliferation Kit (BD Pharmingen), following the manufacturer’s instructions. Briefly, H1299 cells were plated in 6-well plates with 10% FBS medium. After attachment, cells were starved with 0.5% FBS medium overnight and treated with test compounds. Treated cells were harvested, stained with Alexa Fluor^®^ 647 Mouse Anti-H2AX (pS139) antibody and PE Mouse Anti-Cleaved PARP (Asp214) Antibody. The signals were assessed with a CytoFlex flow cytometer (Beckman Coulter Life Sciences).

### 4.16. Statistical analysis

Two tailed Student’s *t* tests were used for data analysis with two groups. One-way analysis of variance (ANOVA) was used for data analysis with no less than three groups followed by Dunnett’s test for comparison between different groups. A P value ≤ 0.05 was considered statistically significant. All analyses were conducted in GraphPad Prism4 (GraphPad Software, Inc.)

## Acknowledgements

The work was partly supported by grants from the National Institute of Health (R01CA193286) to C. Xing, the Harry T. Mangurian Jr. Foundation to C. Xing, College of Pharmacy Frank Duckworth Endowment to C. Xing, University of Florida Health Cancer Center Startup fund to C. Xing, and University of Florida Medicinal Chemistry Mass Spectrometry Support to C. Xing. Y. Wu was supported by a scholarship from the National Science Foundation of China. The funders had no role in study design, data collection and interpretation, or the decision to submit the work for publication. We also acknowledge National Cancer Institute providing PLCO information used in this manuscript. We would like to thank Sreekanth C. Narayanapillai for the help of mouse tissue sample collection. We would like to thank Santanu Hati for characterizing NNAL chirality.

The costs of publication of this article were defrayed in part by the payment of page charges. This article must therefore be hereby marked advertisement in accordance with 18 U.S.C. Section 1734 solely to indicate this fact.

