## Supplemental material for "NNAL, a major metabolite of tobacco-specific carcinogen NNK, promotes lung cancer progression through deactivating LKB1 in an isomer-dependent manner"



Key words: tobacco, lung cancer, NNK, (R)-NNAL, LKB1

**Supplementary Figure 1.** Effect of NNAL exposure on H1975, HCC827 and H460 cell proliferation. *, P<0.05; **, P<0.01; ***, P<0.001. Treatment 6 days.

**Supplementary Figure 2.** Effect of NNAL enantiomers (100 nM) on gemcitabine or cisplatin treatment in H460, H1975 and HCC827. *, P<0.05; **, P<0.01; ***, P<0.001. Treatment 2 days.

**Supplementary Figure 3.** Effect of NNAL enantiomers on CREB activating phosphorylation at Ser133 in H1299, H1975, HCC827, A549 and H460 cells. NNAL treatment 30 min.

**Supplementary Table 1.** Antibodies used for WB.

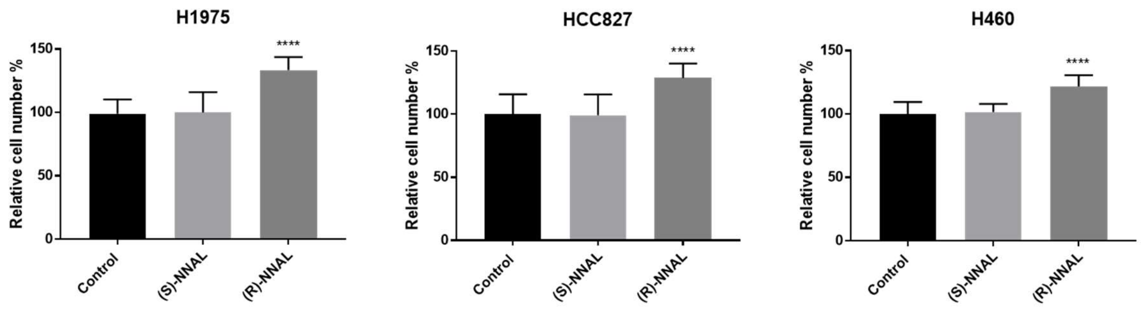
H1975
HCC827
H460
Relative cell number %
150
100
50
0
****
Control
(S)-NNAL
(R)-NNAL

Fig. S1.

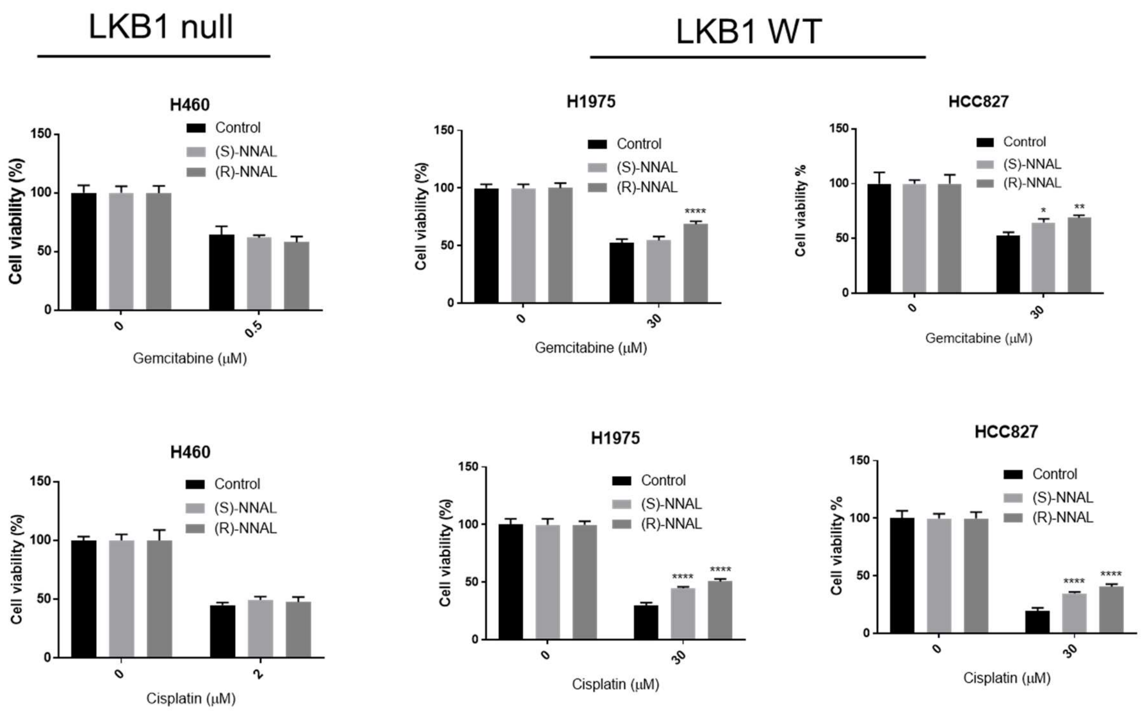
LKB1 null
LKB1 WT
H460
Control
(S)-NNAL
(R)-NNAL
Cell viability (%)
Gemcitabine (μM)
H1975
****
HCC827
*
**
Cell viability %
Cisplatin (μM)
Fig. S2.

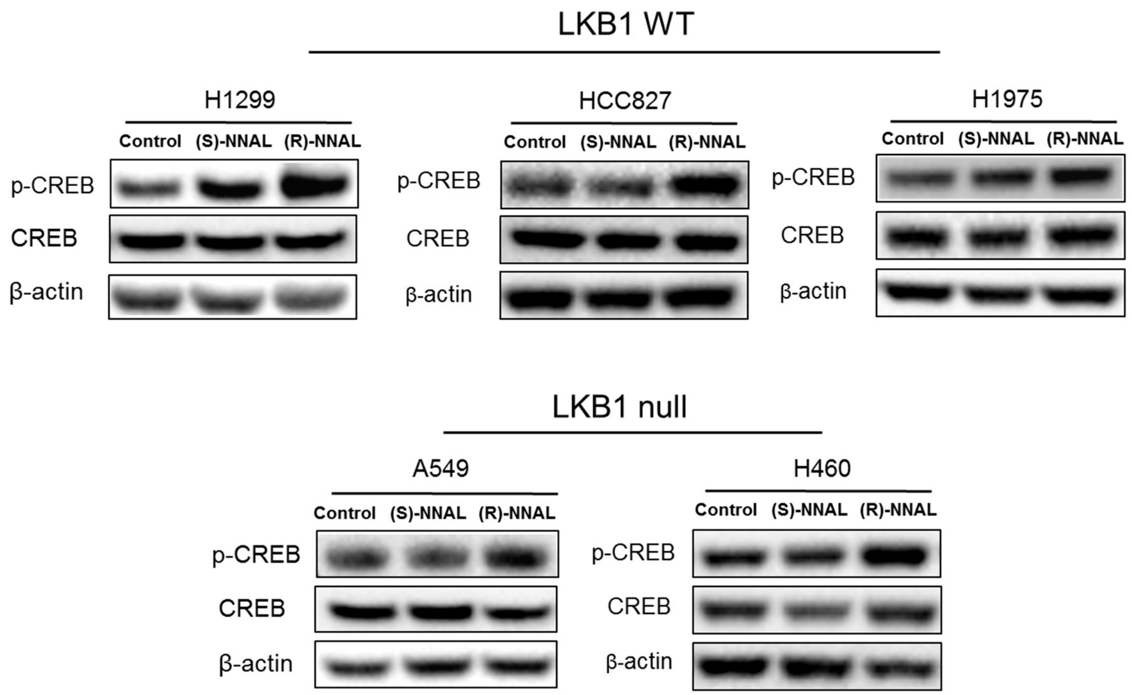

LKB1 WT
H1299
Control
(S)-NNAL
(R)-NNAL
p-CREB
CREB
β-actin
HCC827
Control
(S)-NNAL
(R)-NNAL
p-CREB
CREB
β-actin
H1975
Control
(S)-NNAL
(R)-NNAL
p-CREB
CREB
β-actin
LKB1 null
A549
Control
(S)-NNAL
(R)-NNAL
p-CREB
CREB
β-actin
H460
Control
(S)-NNAL
(R)-NNAL
p-CREB
CREB
β-actin


Fig. S3.

| Antibodies | Company | Catalog number |
|---|---|---|
| LKB1 (D60C5) Rabbit mAb | Cell Signaling Technology | 3047 |
| Phospho-LKB1 (Ser428) (C67A3) Rabbit mAb | Cell Signaling Technology | 3482 |
| CREB (48H2) Rabbit mAb | Cell Signaling Technology | 9197 |
| Phospho-CREB (Ser133) (87G3) Rabbit mAb | Cell Signaling Technology | 9198 |
| AMPKα Antibody | Cell Signaling Technology | 2532 |
| Phospho-AMPKα (Thr172) (40H9) Rabbit mAb | Cell Signaling Technology | 2535 |
| Bcl-2 (124) Mouse mAb | Cell Signaling Technology | 15071 |
| Bim (C34C5) Rabbit mAb | Cell Signaling Technology | 2933 |
| PCNA (PC10) Mouse mAb | Cell Signaling Technology | 2586 |
| Cleaved PARP (Asp214) (D64E10) XP® Rabbit mAb | Cell Signaling Technology | 5625 |
| Phospho-4E-BP1 (Thr37/46) (236B4) Rabbit mAb | Cell Signaling Technology | 2855 |
| 4E-BP1 (53H11) Rabbit mAb | Cell Signaling Technology | 9644 |
| Phospho-mTOR (Ser2448) (D9C2) XP(R) Rabbit mAb | Cell Signaling Technology | 5536 |
| mTOR (7C10) Rabbit mAb | Cell Signaling Technology | 2983 |
| Anti-mouse IgG, HRP-linked Antibody | Cell Signaling Technology | 7076 |
| Anti-rabbit IgG, HRP-linked Antibody | Cell Signaling Technology | 7074 |
| Mouse monoclonal anti-beta-actin | Sigma Aldrich | A2228 |

Table S1.
